# Not by systems alone: replicability assessment of disease expression signals

**DOI:** 10.1101/128439

**Authors:** Sara Ballouz, Max Dörfel, Megan Crow, Jonathan Crain, Laurence Faivre, Catherine E. Keegan, Sophia Kitsiou-Tzeli, Maria Tzetis, Gholson J. Lyon, Jesse Gillis

**Author notes:** Corresponding author: Dr Jesse Gillis, The Stanley Institute for Cognitive Genomics, Cold Spring Harbor Laboratory, Cold Spring Harbor, NY, 11724, USA. Contact Information: JG, GJL, SB, MD, MC, JC, LF, CEK, MT, SKT. Current addresses (Max Dörfel, Jonathan Crain, Gholson J. Lyon).

## Abstract

In characterizing a disease, it is common to search for dysfunctional genes by assaying the transcriptome. The resulting differentially expressed genes are typically assessed for shared features, such as functional annotation or co-expression. While useful, the reliability of these systems methods is hard to evaluate. To better understand shared disease signals, we assess their replicability by first looking at gene-level recurrence and then pathway-level recurrence along with co-expression signals across six pedigrees of a rare homogeneous X-linked disorder, *TAF1* syndrome. We find most differentially expressed genes are not recurrent between pedigrees, making functional enrichment largely distinct in each pedigree. However, we find two highly recurrent “functional outliers” (*CACNA1I* and *IGFBP3*), genes acting atypically with respect to co-expression and therefore absent from a systems-level assessment. We show this occurs in re-analysis of Huntington’s disease, Parkinson’s disease and schizophrenia. Our results suggest a significant role for genes easily missed in systems approaches.

## Introduction

Gene dysfunction in disease is frequently studied through a systems biology framework (Kitano, 2002) in which genes are linked through their shared functional properties and pathways. To assay the networks underpinning a phenotype it is common to look to the transcriptome as a measure of the activity of the genes in the system (Cookson et al., 2009; Dermitzakis, 2008; Jirtle and Skinner, 2007; Lamb et al., 2006). Gene expression changes are detected through differential expression, and then systems-level signals are determined through gene set enrichment (Hosack et al., 2003; Subramanian et al., 2005), network connectivity assessments (Barabási et al., 2010; Greene et al., 2015; Lage et al., 2007), or gene property enrichment (Kircher et al., 2014; Lek et al., 2016). These methods all contextualize genes to known biological properties, functions and pathways. As this step aims to summarize the disease mechanism, it is typically the last step in an analysis, with little direct evaluation for efficacy (Nguyen et al., 2016; Pham et al., 2017; Yu et al., 2017). This reflects the challenge of developing a systematic framework for doing so, particularly where normal heterogeneity of phenotype and genotype may generate joint functional signals of their own. In this work, we seek to assess the replicability of disease and systems biology signals using a meta-analytic approach that exploits multiple pedigrees of a rare and homogeneous disorder, the *TAF1* syndrome.

Genes may share signals for either biological or technical reasons. In gene or protein space, a “systems biology analysis” is used to define or assess these shared signals, with the assumption that the main driver behind the common features is biological (Draghici et al., 2007; Huang et al., 2008). A systems analysis can take the form of enrichment (e.g., Gene Ontology annotation overlaps) or more complex methods (e.g., *k*-nearest neighbors in co-expression networks), but ultimately outputs pathway-level summaries. Signals shared between genes due to technical properties such as sampling biases, confounded study designs, and batch effects can arise as false positives in a systems biology analysis and may be difficult to identify. False positive results can also easily arise due to unknown or uncontrolled biological variation, stemming from difficulties in phenotyping and the genetic heterogeneity of complex disorders (e.g., schizophrenia and autism (Purcell et al., 2014; Sanders et al., 2017)). There, variation in either phenotype or genotype may average away real disease effects and/or generate gene expression variation unrelated to disease (Hansen et al., 2011) (Figure 1). Replication is a central test of which of the two - either disease specific or untargeted variation - has driven the appearance of a characteristic signal across genes. In this work we assess the recurrence of candidate genes and pathways across separately analyzed data-sets. We characterize the degree to which replicate transcriptional signatures occur in groups of genes (e.g., co-expression) versus outliers (genes acting alone). A particular target of our analysis is the *TAF1* syndrome cohort, a rare and well-defined X-linked neurodevelopmental disorder with multiple pedigrees for assessment (see Box 1 for further details).

**Figure 1:**
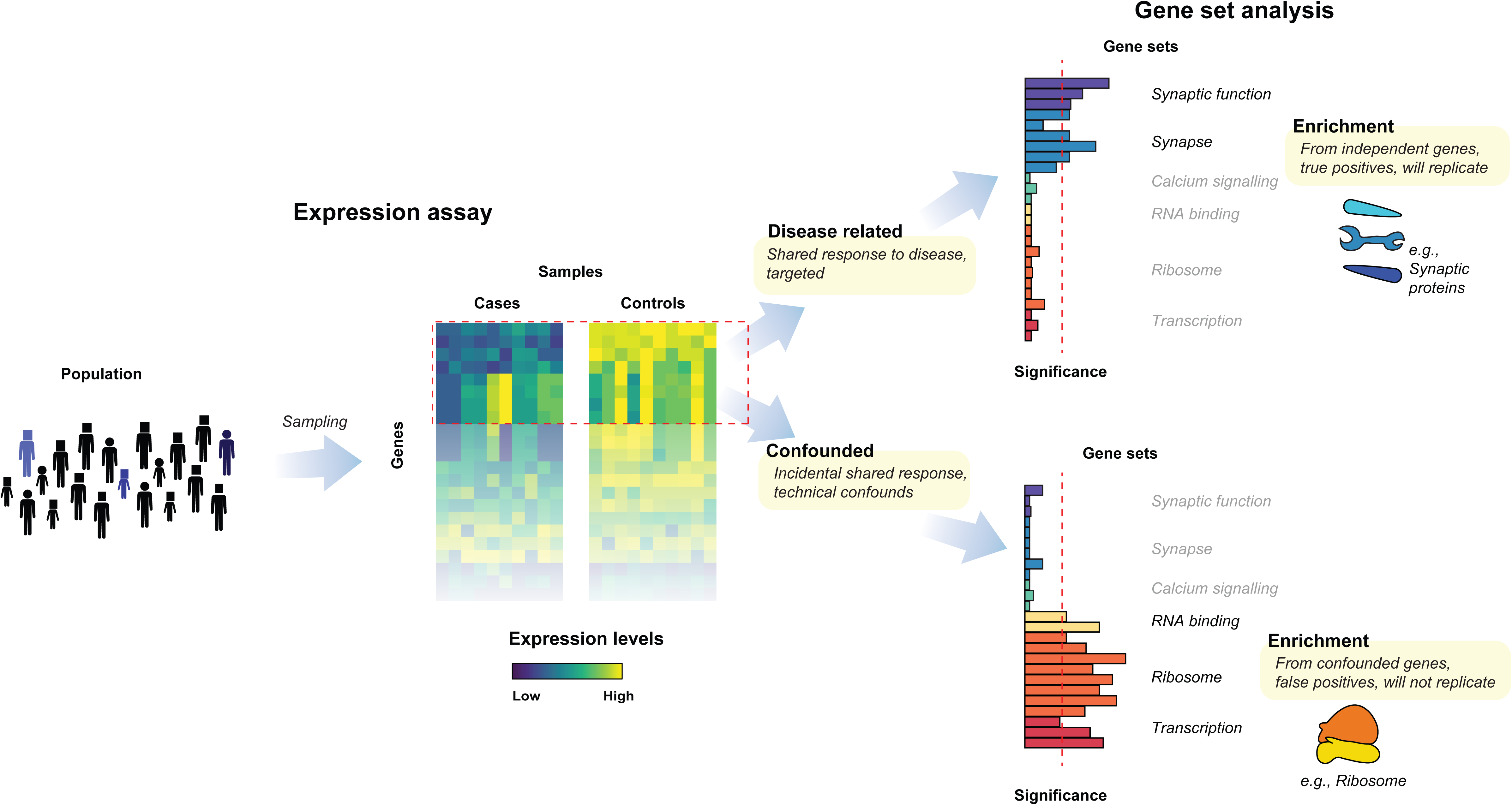
What does a systems biology approach tell us? A systems biology assessment summarizes the properties of genes that are captured in an experiment, but can highlight both true and false positive results. In some cases, genes with a shared function are unlikely to arise in the experiment by chance. A gene set analysis will highlight their shared function (top panel), but not their independent value in identifying the enriched function. In other cases, a set of genes are so closely related both technically and biologically that if one arises, the others are almost certain to do so. A statistical analysis treating the genes as independent (bottom panel) will attach a misleading significance to the shared presence of the genes. These genes and gene sets will be unlikely to replicate in future studies.

#### Box 1 *TAF1* syndrome cohort

To probe in detail the functional gene signals that recur significantly due to disease, we focus a large part of our analysis on *TAF1* syndrome also known as “X-linked syndromic mental retardation-33” (*MRXS33* MIM# 300966 (O’Rawe et al., 2015)), an X-linked recessive neurodevelopmental disorder. *TAF1* syndrome is a rare, penetrant, and overall homogeneous disorder with no known disease mechanism. Genetically, it is defined by mutations in *TAF1* (TATA-Box Binding Protein Associated Factor 1), a key subunit of the general transcription factor TFIID (Louder et al., 2016; Müller et al., 2010). TFIID promotes transcriptional initiation by binding to the core promoters of genes, and recruits other transcription factor subunits that act as co-activators or co-repressors, encoding regulatory specificity (Pijnappel et al., 2013). Other subunits of TFIID are candidate genes in developmental and neurodegenerative diseases (Alazami et al., 2015; Bauer et al., 2004; El-Saafin et al., 2018; Hellman-Aharony et al., 2013), with reduced binding between the subunits playing a role in the pathogenesis. The characteristic phenotypic features of the *TAF1* disorder include global developmental delay, facial dysmorphology, generalized hypotonia, hearing impairments, microcephaly, and a characteristic gluteal crease with a sacral caudal remnant. All documented cases of the disorder affect males, and mostly have arisen *de novo*. In the few known inherited cases, female carriers do not show any features of the disease. This is generally a feature of X-linked disorders, with extreme X-skewed inactivation playing a role in phenotypic variation and protection in females (Migeon, 2007).

Despite being a relatively rare disorder, we have access to multiple pedigrees. In this study, six families were recruited from around the world, mainly of European descent and were between 5-21 years of age. All probands have point mutations in their *TAF1* transcription factor, except for a single CNV case. In three of the pedigrees, the mothers are carriers of the same mutation.

The four properties – global transcriptional impact, characteristic phenotype, genetic homogeneity, and multiple pedigrees - allow us to perform a disease replicability analysis using easily accessible blood transcriptional profiles. We can study each pedigree as a separate differential expression experiment: identifying differentially expressed genes and overrepresented pathways and then assessing these candidate genes and pathways for recurrence across pedigrees.

We are able to look for replicability of disrupted gene expression signatures within the *TAF1* cohort for four main reasons: multiple pedigrees, phenotypic similarity, genetic homogeneity and a plausible mechanism for an impact on expression levels. We describe four classes of signals that can be extracted from disease analyses, reflecting whether signals are shared across families (recurrent/replicable) and across functional sets of genes (joint/disjoint signals). In *TAF1* syndrome, we find recurrence of gene expression change at the gene-level for two plausible candidates, *CACNA1I* and *IGFBP3*. At the systems biology level, we find little replicable enrichment, assessed via gene set enrichment of the Gene Ontology (GO) and via co-expression. Interestingly, the strongest replicable signal appears to arise from genes acting outside of the systems biology framework. We call genes meeting this property “functional outliers”. To see if our analysis is informative in other diseases and study designs, we also assess common heterogeneous neurodegenerative and neuropsychiatric disorders in over a thousand samples (∼1.7k), including 5 Huntington’s disease studies, 15 Parkinson’s disease studies, and 10 schizophrenia studies. Once we extend the analysis to the more heterogeneous brain disorders, we are able to recapitulate known disease mechanisms, which appear as both recurrent joint signals (e.g., immune signaling pathways in schizophrenia), and recurrent functional outliers (e.g., *SNCA* in Parkinson’s). The results from the four disorders studied here highlight the potentially important role of functional outliers, and suggest caution in applying gene-set based methods, such as enrichment or co-expression, in summarizing disease manifestation and mechanism.

## Results

### Replicability design overview

To understand shared disease signals, we assess replicability by testing the recurrence of candidate genes and pathways across separately analyzed differential expression experiments. We classify whether signals are replicated across families or datasets (and call this “recurrent”) and whether the signals involve sets of genes acting together (and call this “joint”). For the *TAF1* syndrome cohort analysis, we performed a family-based differential expression analysis, then tested for gene-level and pathway-level signals through gene set enrichment and a co-expression network modularity analysis (Figure 2A). We then looked across the pedigrees to evaluate where signals arise, summarized in Figure 2B. There are four possibilities we can assess. 1) We may have a joint functional signal, where many genes are part of the same set/module, and it is this module which is replicated across families (i.e., recurrent and co-functional). 2) We may have a recurrent disjoint signal: genes that replicate across families, but do not share common functions with other genes (i.e., recurrent functional outliers). 3) There is also the chance of a non-recurrent but joint signal: genes contribute to a shared signal, but uniquely within a family. 4) And finally, there is also an entirely disjoint signal, where we see one-off genes that are most likely false positives. We test all possible outcomes by measuring recurrence of genes, gene set enrichment and co-expression modularity (Figure 2C). We go through the results in the following sections.

**Figure 2:**
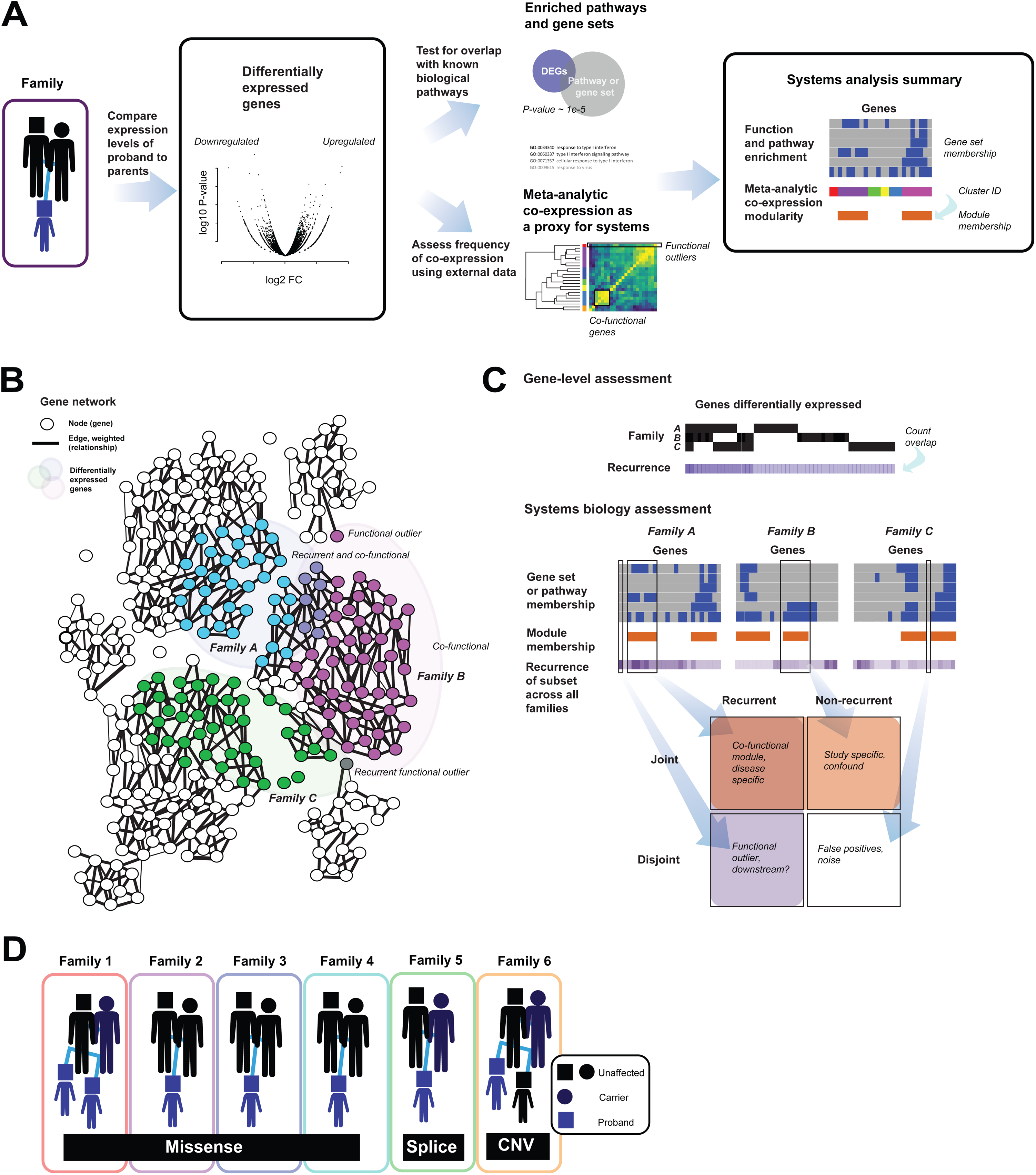
Disease expression analysis schematic. (A) We calculate expression fold change between probands and parents and pick out the top 100 up-and down-regulated genes (increased and decreased expression, respectively). We test for joint functional properties through gene set enrichment and co-expression modularity. (B) Given multiple pedigrees/families of the disorder, we can piece together whether disease signals are recurrent across families or non-recurrent, and due to multiple genes (joint) or independent genes (disjoint). (C) This can be done by assessing recurrence at the gene and pathway levels. (D) The *TAF1* syndrome cohort pedigrees used in this analysis. Four cases have missense mutations, one case a splice site mutation, and the last case a CNV duplication. Three of the mothers are carriers with no distinguishing characteristics.

### Transcriptional replicability across the *TAF1* cohort occurs at the gene level

Since *TAF1* is a transcription factor, we first wished to see if there was a common disease signature at the expression level. We identified differentially expressed genes (DEGs) using a family-based differential expression (DE) analysis (see **STAR Methods**). We saw only moderate overlaps of the differentially expressed genes between each of the pedigrees tested (Figure 3A) for both upregulated and downregulated genes. There were at most 26 (out of 100) genes in common between Family 3 and Family 5 (Figure 3B, p∼1e-40). Even though few genes overlapped when assessed pairwise, there were a number of differentially expressed genes that were recurrently DE across families (significant if in at least three families FDR<0.05), with a modest number significantly recurrent. We find four genes recurrently upregulated (*ISG15, RN7SK, FFAR3* and *IGLV1-44*), and 14 downregulated (*C1QA, CFH, RPS7, SNRPG, LSM3, RPS3A, IGFBP3, RPL7, KLRB1, OLFM4, PRSS30P, KIR2DS4, S100B* and *CACNA1I* Figure 3C, see **Table S4**). These results suggest that even though a large fraction of the differentially expressed genes are unique to each pedigree, a recurrent gene transcriptional disease signature is present within the data. To test the dependence of these results on the DE threshold (top 100 genes), we repeated the recurrence using different DE thresholds (Figure 3D). As the change in the number of the DEGs called had an influence on the significance of recurrence, we see peaks and troughs of gene recurrence corresponding to the change in adjusted p-values. We found similar numbers of significantly recurrent genes within these ranges, with cut-offs between 50 and 200 genes most informative. We use the top 100 DEGs for the remainder of the analyses.

**Figure 3:**
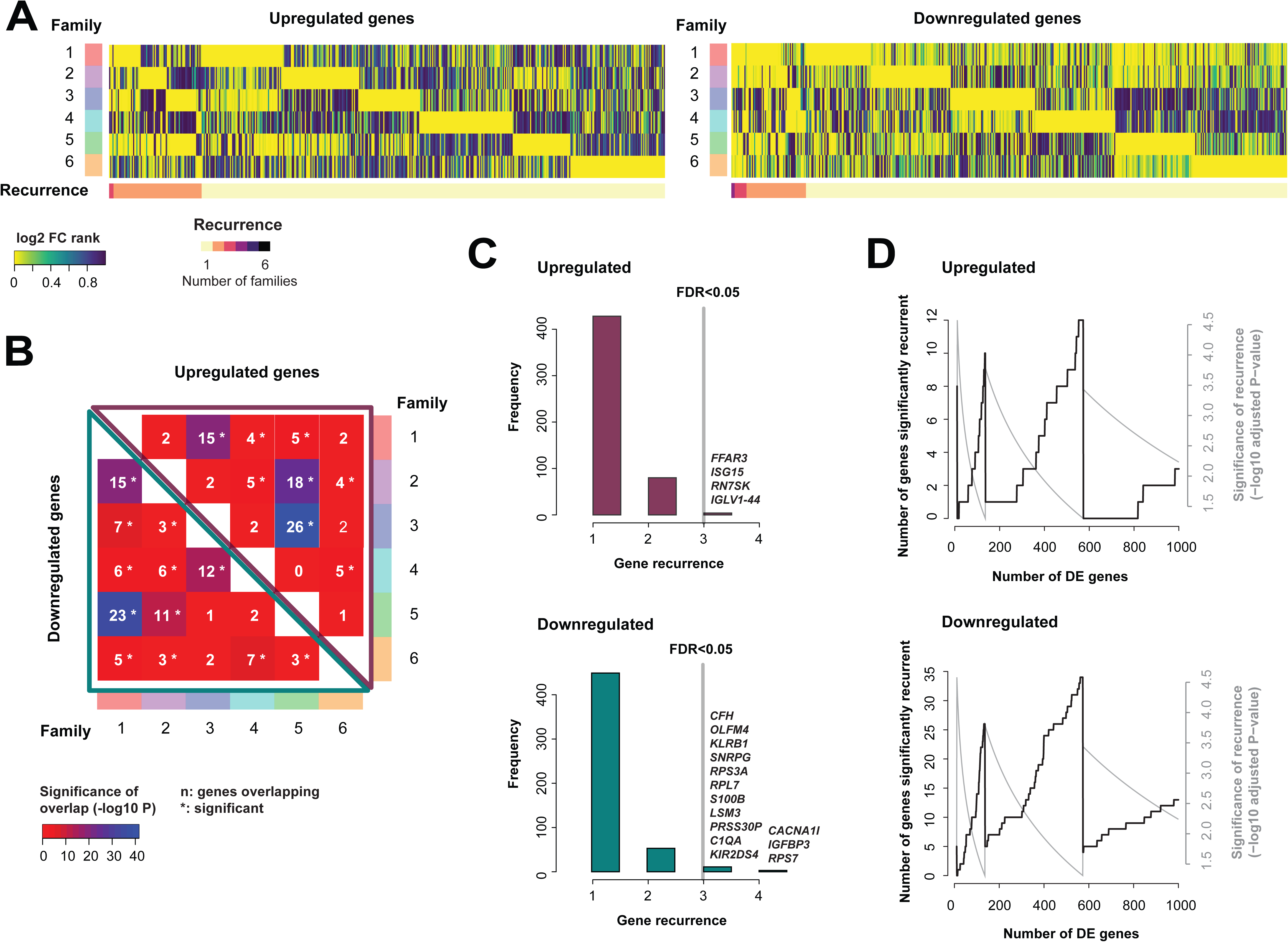
Disease expression analysis with a family-based approach. (A) The expression fold change for each gene is calculated within each family (top 100 up and down regulated genes are shown). (B) Overlaps in DE gene sets between the individual families (numbers in boxes), and the significance of this overlap (colored corresponding to - log10 P-value of the hypergeometric test). Overlaps are mostly small. (C) The replicable genes are those that are recurrent across families. The recurrence distributions for both up-and downregulated genes across the 6 families are shown. Using the binomial test, we find that genes recurring 3 or more times are significant (FDR<0.05). These genes are listed, with 4 up-and 14 downregulated genes significantly recurrent. (D) Robustness assessment of the DE threshold. The plot shows the number of recurrent genes as a function of the number of differentially expressed genes and the significance of the recurrence in grey.

### Gene set enrichment does not identify specific disease mechanisms

While only a modest number of genes overlapped across the pedigrees, it is possible that non-recurrent genes still provide a shared disease signal, with variability in the exact genes identified due to technical limitations. To assess this possibility, we perform gene set enrichment in two ways. First, we test differentially expressed gene lists from each pedigree independently, and then measure the recurrence of the enriched pathways across the studies, as a parallel to the gene recurrence assessment. We then check for significantly recurrent pathways. Second, we test for enrichment of the recurrent genes themselves. We performed gene set enrichment using a subset of the Gene Ontology (GO slim) and found few pathways significantly enriched on a per family basis (FDR<0.05, Figure 4A), with the exception of Family 5, which had 22 downregulated pathways. Then, to assess the disease significance of the enrichment signal, we calculated the similarity and significance of recurrence of these pathways across the families. As in the case of gene recurrence, we expect the disease signal (here the enrichment signal) to replicate across the cohort if it is disease linked. Of the significantly enriched pathways, at most three were significantly upregulated and recurrent across the families (Figure 4B and **Table S5**). The pathways significantly recurrent were larger than average, implying broad properties (e.g., GO:0002376 immune system process, GO:0006950 response to stress and GO:0007165 signal transduction, **Figure S3**A) with different genes contributing to the enrichment within each of the families (Figure 4C). While pathways were significantly recurrent across family-specific upregulated genes, the few recurrent upregulated genes do not show enrichment. In contrast, we found no downregulated pathways as significantly recurrent (Figure 4D) despite the gene recurrence and family-specific enrichment. However, the recurrent downregulated genes themselves (Figure 4E) are enriched for six GO groups related to ribosomal functions, including RNA binding, translation, and protein targeting. The discrepancy between the pathways enriched with the recurrent genes and the pathways significantly recurrent suggests that the weak enrichment signals are not specific to the disorder.

**Figure 4:**
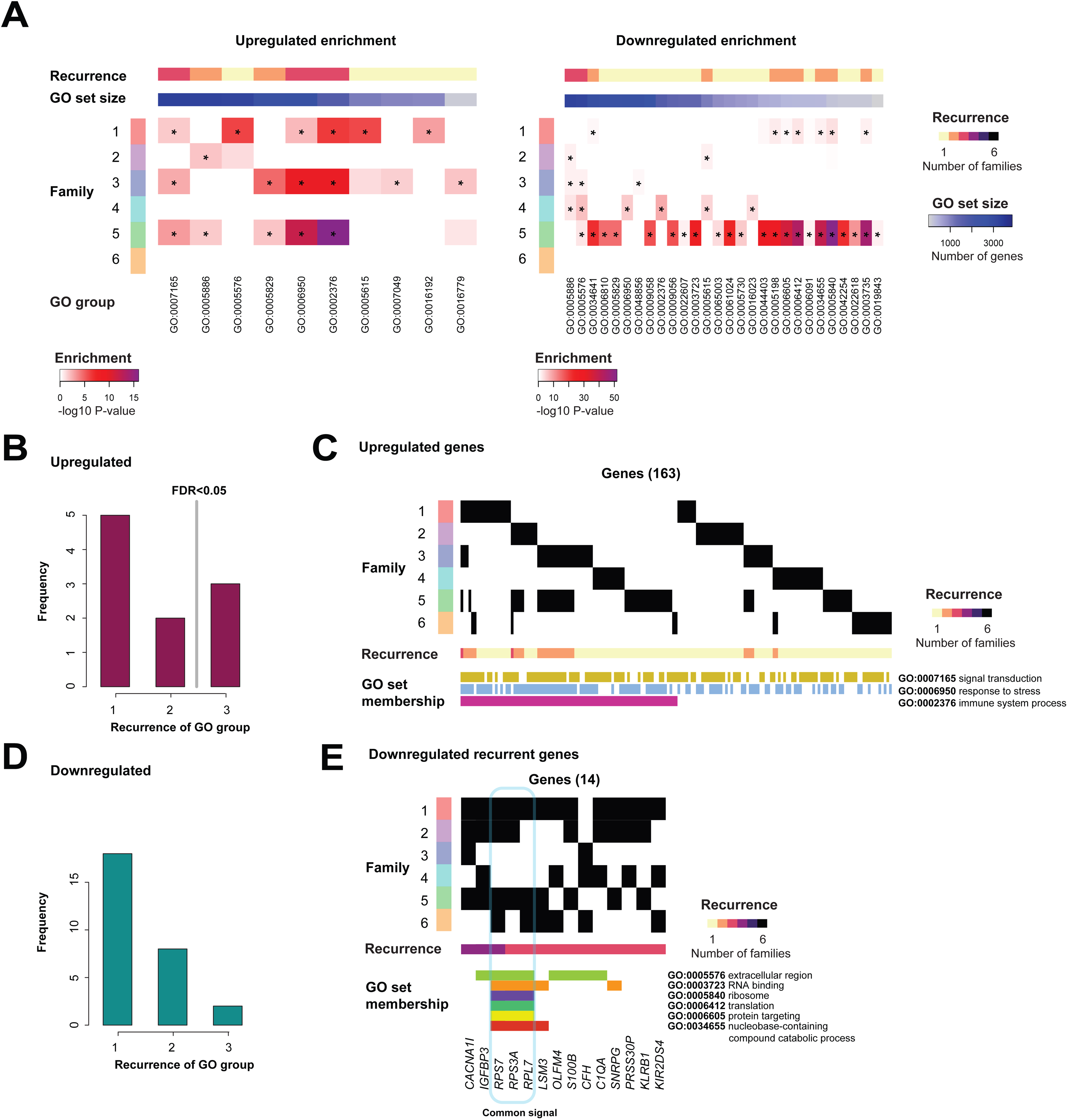
Gene set enrichment assessment of *TAF1* cohort. (A) Top GO enrichment results for each family for up-, and downregulated genes. Significant terms (FDR < 0.05) are highlighted with an *. (B) The frequency and significance of recurrence of each GO term is plotted for the upregulated genes. (C) Gene-GO membership matrix for the upregulated genes. Each column is a gene, and each row a family. The colored bars below highlight the GO terms that these genes belong to. The signal associated with the recurrent GO terms is distributed across different genes, shown by low overlap across the families. (D) There are no significantly recurrent pathways with the downregulated genes. (E) However, the recurrent genes themselves are enriched for ribosomal pathways (p-adjusted<0.05), as shown in the gene-GO membership matrix. The three RP* genes seem to drive almost all the signal.

### Top recurrent genes do not appear to act within a systems biology framework

As we are limited by gene set annotations, the weak enrichment may be due to our lack of complete pathway knowledge (Thomas, 2017). Therefore, to test for joint signal with a different but comprehensive data modality, we looked to co-expression. Genes that are co-expressed are known to share functions, are co-regulated, or are parts of known pathways (Gaiteri et al., 2014). Unlike most curated or inferred gene annotations, co-expression can be assessed genome-wide. We used the gene pair co-expression frequency from a wide corpus of data (external to this study) as our co-functionality measure (see **STAR Methods**). For each family’s set of differentially expressed genes, we found gene co-expression blocks comprising more than two thirds of these genes (**Figure S4**A). We show an example of this for the top 100 down-regulated genes from Family 1 in Figure 5A.

**Figure 5:**
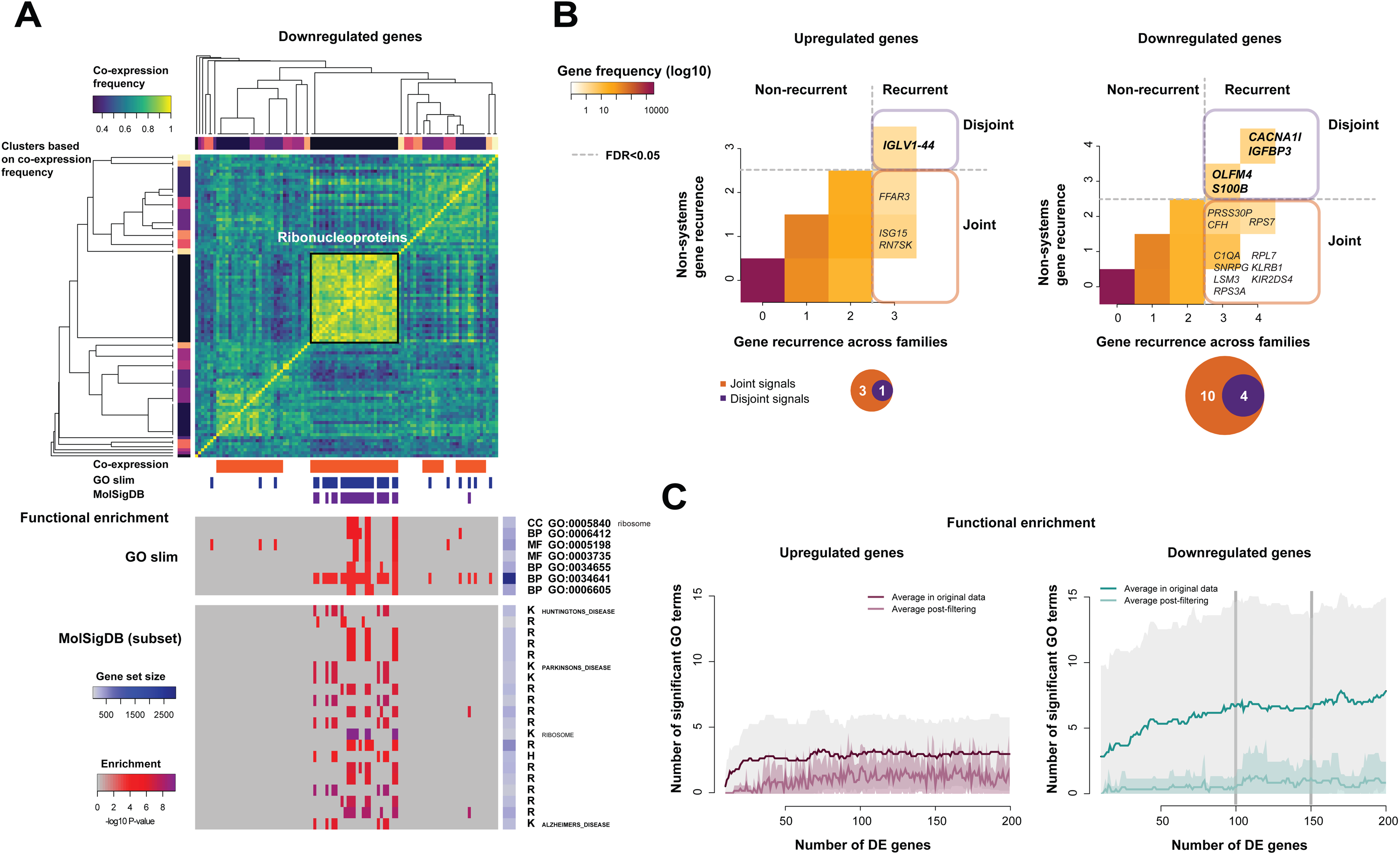
Co-expression of differentially expressed genes generates enrichment. (A) As an example from family 1, we show the co-expression frequency sub-network as a heatmap, where genes showing decreased expression show co-expression. Co-expression blocks define modules as determined by the clustering (see rows). The modules are enriched for particular genes, mainly ribonucleoproteins. Performing a gene set enrichment analysis on these genes (Fisher’s exact test on GO groups), genes (rows) that generate the enrichment (columns are enriched GO terms) almost exclusively overlap with the co-expression blocks. The prominent pathways are ribosome related. (B) The significantly recurrent genes can be divided into those present within co-expression modules (joint) and those not (disjoint). The genes in bold are the functional outliers and the venn diagrams summarizes the number of genes in each category. (C) If we look at the enrichment of these DE gene sets (pre-filtering dark line +/-SD shadow), we see that filtering off the modules removes all but a few significant terms (lighter line, +/-SD shadow).

We then asked where the recurrent genes sit in relation to the co-expression modules. Interestingly, we found the outlier or disjoint genes (those not in the large modules) were frequently among the most recurrent genes across the families (Figure 5B). Almost a third of the disease signature was not within common co-expression or functions, but rather appeared within very small modules or as outliers. For functional characterization to usefully summarize the DE list, candidate genes should be enriched within pathways. This was not true of *TAF1* disorder candidate genes. In particular, we find that the top downregulated genes *CACNA1I* and *IGFBP3* are not within modules. Once we perform a gene set enrichment analysis on the genes excluding those in modules, nearly all of the enrichment signal is lost, highlighting that the most recurrent candidates act outside of a common joint signal. If this is similarly true of the disease mechanism, then a systems-style analysis will fail to discover the disease signal.

### Recurrence of genes and pathways in other disorders

The *TAF1* cohort is interesting in part for being a rare disorder with a penetrant and distinct phenotype. In order to assess the role of functional outliers more broadly, we looked to three other disorders with substantial transcriptomic data and varying degrees of genetic heterogeneity. We focused on Huntington’s disease (HD), Parkinson’s disease (PD), and schizophrenia (SCZ). All studies used are listed in **Table S3**.

Huntington’s disease is an inherited neurodegenerative disorder, characterized by the progressive degeneration of cells in the brain (primarily the striatum) and is associated with impaired movements, decline in cognitive abilities and depression (Bates et al., 2015). Similar to *TAF1* syndrome, HD is a monogenic disorder. The disease is caused by an expansion of the CAG repeats in the Huntingtin gene (*HTT*), which is believed to be toxic to other proteins once mutated. The exact functions of *HTT* are still unclear, along with the mechanisms (Bates, 2005). To test the possibility of a joint functional signal, we assessed five expression studies of HD, following the evaluative approach we took with the *TAF1* cohort. Interestingly, we found both joint and disjoint signals (Figure 6A). A total of 18 upregulated genes were significantly recurrent, with *ANGPT1* (Angiopoietin 1) recurrent in four of the five studies. This gene plays an important role in vascular development and angiogenesis. Along with other recurrent genes, such as *ANG* (angiogenin), *KCNE4* and *SLC14A1*, there seems to be an associated cardiovascular phenotype. Heart disease is comorbid with Huntington’s, and these gene candidates suggest a link to cardiac development. Additionally, we found six genes recurrently downregulated in at least three of the five studies; these include synaptic genes (*SV2C, NRGN*, and *HTR2C*) and a calcium channel related to modulation of firing in neurons (*CACNA1E*). We found that all the recurrent downregulated genes were part of modules, thus showing a strong joint functional signal. And in the upregulated genes, *ANG* is a functional outlier, while all of the other recurrent genes were in shared functions. These results suggest a stronger joint functional signal than in the *TAF1* disorder. All recurrent genes are listed in **Table S6**.

**Figure 6:**
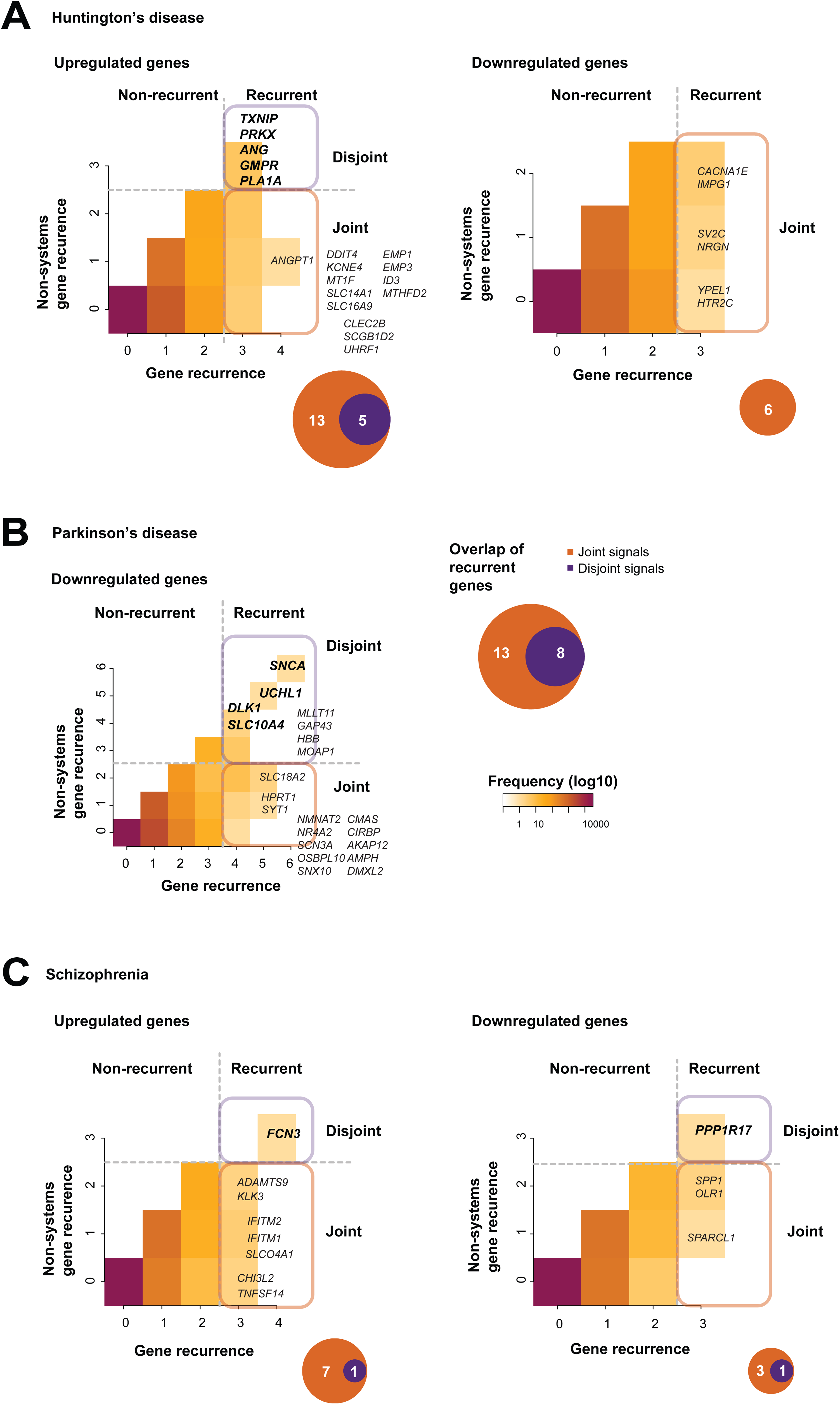
Differential expression meta-analysis in three other disorders. (A) Recurrence of genes in Huntington’s disease (HD), (B) Parkinson’s disease (PD) and (C) schizophrenia (SCZ), and whether they occur in groups (joint) or not (disjoint). The venn diagrams summarize the number of recurrent genes and their joint or disjoint designation.

Parkinson’s disease is a progressive neurodegenerative disorder, characterized by the loss of dopaminergic neurons leading to decreased motor function. Unlike the *TAF1* cohort, Parkinson’s has multiple genes implicated, each with different onset stages (e.g., *LRRK, SNCA, PRKN, FBXO7, PARK7,* and *PINK1*) (Poewe et al., 2017), increasing the genetic heterogeneity of the disorder and data. We collected 15 differential expression gene lists and repeated our analysis. We find a subset of significantly recurrent downregulated genes but no upregulated genes as significant (Figure 6B). The most recurrently downregulated gene is *SNCA* (alpha-synuclein), a well-known Parkinson’s disease gene (Siddiqui et al., 2016) which recurs six times (FDR < 0.05). We did not purposely select for studies with variants in this gene when selecting the studies for the meta-analysis, but could confirm it was the genetic cause in a few of the studies. Another top candidate was *SYT1* (synaptotagmin 1) which recurred in five of the 15 studies (FDR < 0.05). Synaptotagmins are known to be involved in neurodegeneration (Glavan et al., 2009), and *SYT11* (Sesar et al., 2016) has been associated with the disorder through its interactions with *PARK7*. Other genes that are significantly recurrent include *SLC18A2* (linked to PD (Lohr and Miller, 2014)), *UCHL1* (mutations may be associated with PD (Healy et al., 2006; Maraganore et al., 2004)), *DLK1* and *SLC10A4*. A majority of the recurrent genes were present in co-expressed modules, but the joint signal was less predominate than in Huntington’s disease. All recurrent genes are listed in **Table S7** and enriched pathways in **Table S8**.

Our final use case was on schizophrenia, a neuropsychiatric disorder characterized by abnormal social behavior and psychosis, along with other cognitive impairments (Kahn et al., 2015). The disorder has strong environmental and genetic components (Gejman et al., 2010; Rees et al., 2015), with many genes increasing risk. The risk alleles are also shared amongst many other neuropsychiatric phenotypes, making this a hard disorder to classify both genetically and phenotypically, and thus difficult to characterize molecularly. We assessed 10 disease expression studies and found both up-and downregulated transcriptional signatures (Figure 6C), but these recurred in three or four studies at most. Consistent with the genetic and phenotypic heterogeneity of the disorder, we find that recurrent genes have both joint and disjoint signals. Of the genes upregulated, we find recurrence of the *FCN3* gene (Ficolin 3), a recognition molecule in the lectin pathway of the complement system (Garred et al., 2009; Mayilyan, 2012). This is of interest in schizophrenia as it potentially interacts with C4, a known risk allele (Sekar et al., 2016). The remaining recurrent genes were enriched for an inflammatory signature, also recapitulating known schizophrenia etiology. Among recurrent downregulated genes is the protein phosphatase inhibitor *PPP1R17* which is primarily expressed in Purkinje cells in the cortex of the cerebellum, a brain region which may play a role in the disorder (Maloku et al., 2010). Interestingly, both *FCN3* and *PPP1R17* were functional outliers, with the other recurrent genes showing joint functional signals. All recurrent genes are listed in **Table S9** and enriched pathways in **Table S10**. Overall, across all the disorders, the joint signals via co-expression were much stronger than in the *TAF1* syndrome cohort, but there were important functional outlier candidates within each disease.

## Discussion

The main contribution of this work is a rigorous analysis of replicability of functional signals implicated in disease through expression analysis. We describe four classes of signals that can be extracted from disease analyses, reflecting whether signals are shared across families or studies (recurrent) and/or across functional sets of genes (joint). These are: 1) recurrent joint signals, 2) recurrent but disjoint, 3) joint but not recurrent 4) not recurrent and disjoint. Our evaluation of the rare *TAF1* syndrome and three common disorders highlighted an important feature of disease that has been overlooked in the systems paradigm: the role of functional outliers, classified as recurrent but with no joint or shared function.

In the *TAF1* cohort, we believe that the recurrent disjoint genes are the disease signal for reasons within and outside the present analysis. Of the candidates we found, *CACNA1I* and *IGFBP3* had the strongest recurrent signal. *CACNA1I* is a calcium channel subunit, and mutations in calcium channels are known to have similar phenotypes to the cohort here, including intellectual disability, autism and dystonia (Fukai et al., 2016; Lu et al., 2012). In addition, there is a *TAF1* binding site upstream of the gene (Wang et al., 2012). Most convincing is that this gene has been implicated recurrently in other brain disorders; it is the only recurrent missense *de novo* in schizophrenia studies (Gulsuner et al., 2013), one of the few overlapping candidates between schizophrenia and autism. The other recurrent candidate was the insulin-like growth factor-binding protein 3 (*IGFBP3)*. Downregulation of this gene has been recently implicated in a developmental disorder with a behavioral and cognitive phenotype (Perez et al., 2018). Despite these fairly convincing properties, it remains to be seen whether the candidate genes are specific to the *TAF1* cohort, or rather might reflect a more general signature of developmental disorders.

Consistent with the view that functional outliers and aberrantly expressed genes may play a particularly strong role in diseases with rare genotypic variation, we do not find them as the strongest signal in the other disorders. In our meta-analysis, we focused on disorders with varying genetic architecture and those also well powered for assessment. In each case, the meta-analysis was powered to call individual genes as recurrent, identifying both known and novel candidate disease genes. This was not dramatically more than those found assessing across individual families within the *TAF1* cohort, most likely due to the heterogeneity of the other disorders or their study designs. Within each disorder, we observed clear joint functional signals present through the co-expression analysis, but to varying degrees (e.g., strong immune signals in schizophrenia). Interestingly, despite the greater role of joint functional signals within these disorders, plausible functional outliers exist for each, most notably in the case of Parkinson’s disease where known disease genes, such as *SNCA*, appear to be acting outside of their typical behavior as evaluated from co-expression.

Characterizing whether or not genes exhibit expected shared behavior bears strongly on the subject of disease mechanisms. In the transcriptomic analysis of rare disorders, a joint disruption is almost always assumed for disease. Yet, there is potential for unbuffered and uncharacteristic expression changes in a few single genes, particularly when the assumed pathogenic variant is regulatory or the disorder is monogenic (Cummings et al., 2017; Fresard et al., 2018; Kremer et al., 2017). In this case, genes that are no longer under regulatory control, or those that have gained regulation, will act out of place (Zeng et al., 2015; Zhao et al., 2016). These genes could be far downstream in a pathway or cascade, and thus not impact other pathway members directly or immediately. In general, enrichment and other systems biology analyses will miss these single genes that serve as unique bottlenecks. Our results suggest that filtering based on enrichment will systematically remove interesting candidates. This does not mean that systems-style analyses should not be conducted, rather we suggest that they be treated as a tool to classify which genes are operating within a group and which are not. With time, strong candidates in either category may be identifiable and provide valuable and distinct information about disease mechanisms.

## Acknowledgments

The authors would like to thank the families for participating in the study. The authors would like to thank Sanja Rogic and Paul Pavlidis for their assistance with the Gemma RNA-seq data. The authors would also like to thank the following groups for their samples: Micheil Innes from the Department of Medical Genetics and Alberta Children’s Hospital Research Institute and Rosemarie Smith from the Department of Pediatrics at the Barbara Bush Children’s Hospital. This work was supported by a gift from T. and V. Stanley and a grant from the Collaborative Center for X-linked Dystonia Parkinsonism (CCXDP).

## Author Contributions

JG and GJL conceived the project. JG and SB designed experiments. SB performed computational experiments. MD and JC performed wet-lab experiments. SB and JG wrote the manuscript. GJL arranged for blood donation from which RNA was isolated. LF, CEK, MT, and SKT provided blood samples. JG, SB, MD, GJL, MC interpreted data and edited text. All authors read and approved the final manuscript.

All authors read and approved the final manuscript

## STAR Methods

### CONTACT FOR REAGENT AND RESOURCE SHARING

Further information and requests for resources should be directed to and will be fulfilled by the Lead Contact, Jesse Gillis.

### EXPERIMENTAL MODEL AND SUBJECT DETAILS

#### The TAF1 syndrome cohort

We assessed 6 pedigrees of a genetically and phenotypically homogeneous X-linked *TAF1* syndrome cohorts. The original cohort was assembled by O’Rawe et al. (O’Rawe et al., 2015), which included 11 pedigrees from around the world. The probands are male, between 5-21 years of age, have intellectual disability, distinct facial dysmorphology, general hypotonia, hearing impairments, and a characteristic intergluteal crease. Of the 6 pedigrees we studied, all probands had a point mutation in their *TAF1* transcription factor, except for a single CNV case with a duplication of a ∼0.42 Mb region at Xq13.1 that includes *TAF1* and other genes.

### METHOD DETAILS

#### RNA-sequencing and processing

Blood was collected in PAXgene Blood RNA tubes and the RNA was isolated with the PAXgene Blood RNA kit (QIAGEN) according to the manufacturer’s recommendations. The RNA was quantified using NanoDrop. To increase downstream sensitivity, globin mRNA was depleted from the samples using the GLOBINclear Kit (Life Technologies). Briefly, RNA was precipitated with ammonium acetate, washed and resuspended in 14 µl TE (10 mM Tris-HCl pH 8, 1 mM EDTA). Subsequently, for each sample 1.1 µg RNA were hybridized with the provided streptavidin beads and purified. To control for variation in RNA expression data, 1 µl of a 1:100 dilution of ERCC RNA Spike-In control (Thermo Fisher) was added to 1 µg RNA and libraries generated according to the TruSeq Stranded mRNA Library Kit-v2 (Illumina) with the index primers as indicated in

**Table** S1. Quality control of the generated libraries was performed on a Bioanalyzer High Sensitivity DNA chip (Agilent) and the concentration was measured using Qubit dsDNA HS Assay (Life Technologies). To eliminate primer dimers in the libraries, additional purifications were performed using the Agencourt AMPure XP system (Beckman Coulter). The libraries were pooled to 2-10 nM total concentration and sequenced on an Illumina NextSeq 500, PE100, mid output. Libraries were generated independently for each family and family-pools multiplexed and sequenced on separate lanes. ERCC spike-ins included in the preparation were not used for normalization, but rather as a measure of quality control. Families 2, 3 and 4 showed the lowest variation in the ERCCs between family members, while family 5 and 6 had higher technical noise (**Figure S2**). Reads were filtered for QC and artifacts using the fastX toolbox, and then the reads were paired up using an adapted python script (https://github.com/enormandeau/Scripts/blob/master/fastqCombinePairedEnd.py). The reads were aligned to the genome (GRCh38, GENCODE v22 (Harrow et al., 2012)) using STAR (2.4.2a)(Dobin et al., 2012).

#### Differential expression analysis

We calculated fold change between parents and probands for the differential expression analysis. We first calculated the CPM (counts per million) for each individual, and then took the average CPM for the parents and compared it the CPM of the proband. Fold change was defined as the log2 of the ratio of these values after adding a pseudocount of 1. We exploited within-family variance to detect noisy genes, removing genes that showed strong differential expression between the parents (i.e., top 100 up-regulated and top 100 down-regulated genes). After removing these highly variable genes, top up-and down-regulated genes were defined based on ranked fold change. We assessed each family in a separate batch (library preparation and sequencing run), holding technical variation constant in each family and independent across families, so that gene-level recurrence is not expected to differ from the null. By way of analogy, our experimental design resembles the analysis of *de novo* variants in DNA analyses, in which as many factors as possible are held constant in the control group for the proband. The use of unaffected family members as controls provides the closest possible genetic and environmental match for the probands, constraining variability of known importance for expression analysis(Raser and O’Shea, 2005). Although each family-specific analysis is confounded with age and sex, we anticipate that genes detected as differentially expressed that are due to these overlaps can be assessed and identified directly, as these are well-powered properties in many previous studies.

#### Common co-expression frequency network

Human RNA-seq expression data was downloaded from Gemma (Zoubarev et al., 2012). From the total collection of approximately 300 human experiments, we selected 75 expression experiments (3,653 samples) that we could ascertain derived from tissues and not cell lines (listed in **Table S2**). For each experiment, we consolidated our list of genes/transcripts to the ∼30K genes with Entrez gene identifiers, and did not limit either expression level or occurrence of expression. For each experiment with at least 10 samples, we generated a co-expression network by calculating Spearman’s correlation coefficients between every gene pair (Ballouz et al., 2015) and calculated the frequency that a pair of genes was positively co-expressed (Spearman’s correlation coefficient r_s_>0). We used this tally network as a measure of the frequency of common co-expression of the gene pairs. The more observations with a positive correlation, the more commonly co-expressed the pairs are (see **Figure S1**).

#### Gene set enrichment

To calculate gene set enrichment of the differentially expressed genes, we used an in-house gene set enrichment R script based on the hypergeometric test. For each gene set, we calculated the significance of the overlap of the differentially expressed genes with that set, correcting for multiple tests with Benjamini-Hochberg (FDR, p.adjust in R). We used an in-house parsed version of GO (downloaded July 2015). We report GO slim (filtered to 132 GO groups) results primarily in the text but also assess a subset of GO based on gene set size to remove redundancy (10-100 genes per group, 4605 GO terms). In addition, we assess enrichment using four gene set lists from MSigDB (v6, 1127 gene sets, HALLMARK, KEGG, REACTOME and BIOCARTA).

#### Co-expression module and outlier detection

In a set of differentially expressed genes, we defined co-expression modules as highly co-expressed genes, seen as blocks in a co-expression network when clustered and represented as a matrix. Genes that did not cluster or clustered weakly were considered outliers. These genes potentially have stronger co-expression links with other genes outside of the gene set, but are not as well-linked within the subset. To identify these two classes in our list of differentially expressed genes (DEGs), we first extracted the sub-network of the differentially expressed genes from the co-expression frequency network. Then, thresholding on the median co-expression value of the co-expression network, we used this binary network as distance matrix, and performed hierarchical clustering of the genes. This clustering returned a dendrogram of genes that are closer in distance, and we used this dendrogram to define modules within the data. We used the R dynamicTreeCut (Langfelder et al., 2008) package to select modules within the data with a cut height 0.995. We used these clusters to define our co-expression modules, where clusters with more than five genes were labelled as modules and those smaller as co-expression outliers.

#### Recurrence analysis

To test for replicability of the disease signal, we measured differentially expressed gene recurrences and the significance of recurrence as the probability of observing the differentially expressed genes across all the pedigrees. We first calculated the significance of the pairwise overlap between families using Fisher’s exact test (phyper in R). We calculated the significance of recurrence of the differentially expressed genes using the binomial test (pbinom in R), and then corrected for multiple tests using Benjamini-Hochberg (FDR, p.adjust in R). After we filter on differentially expressed genes within modules, we recalculate the significance of recurrence, this time with a permutation test to obtain an FDR. Similarly, we use a permutation test to calculate significance of recurrence of the pathways from the gene set enrichment assessment.

#### Meta-analysis datasets of neurodegenerative and neuropsychiatric disorders

We sought to repeat the systems biology evaluations in other disorders. We collected the reported differentially expressed genes in Parkinson’s disease, Huntington’s disease and schizophrenia studies. The majority of the studies were collected from recent review articles (Genevie et al., 2018; Li and Teng, 2015), and from a search within the Gemma (Zoubarev et al., 2012) database for “parkinson’s disease”, “huntington’s disease” and “schizophrenia”, respectively (listed in **Table S3**). We downloaded fold changes, p-values and adjusted p-values. We removed studies from our analysis where we either could not assess the direction of the differential expression and where no genes passed significance based on log2 fold changes (log2FC|>1) and adjusted P-values (q<0.05).

### QUANTIFICATION AND STATISTICAL ANALYSIS

All statistical analyses were done in R. Significance was defined as an FDR of 0.05 for all statistical tests.

### DATA AND SOFTWARE AVAILABILITY

All R code, scripts and network data is available for download from our github repository (https://github.com/sarbal/redBlocks). The RNA-seq data has been deposited in GEO/SRA under accession number **GSE84891**. All other software used in this analysis is freely available and has been listed in the key resources table.

## KEY RESOURCES TABLE

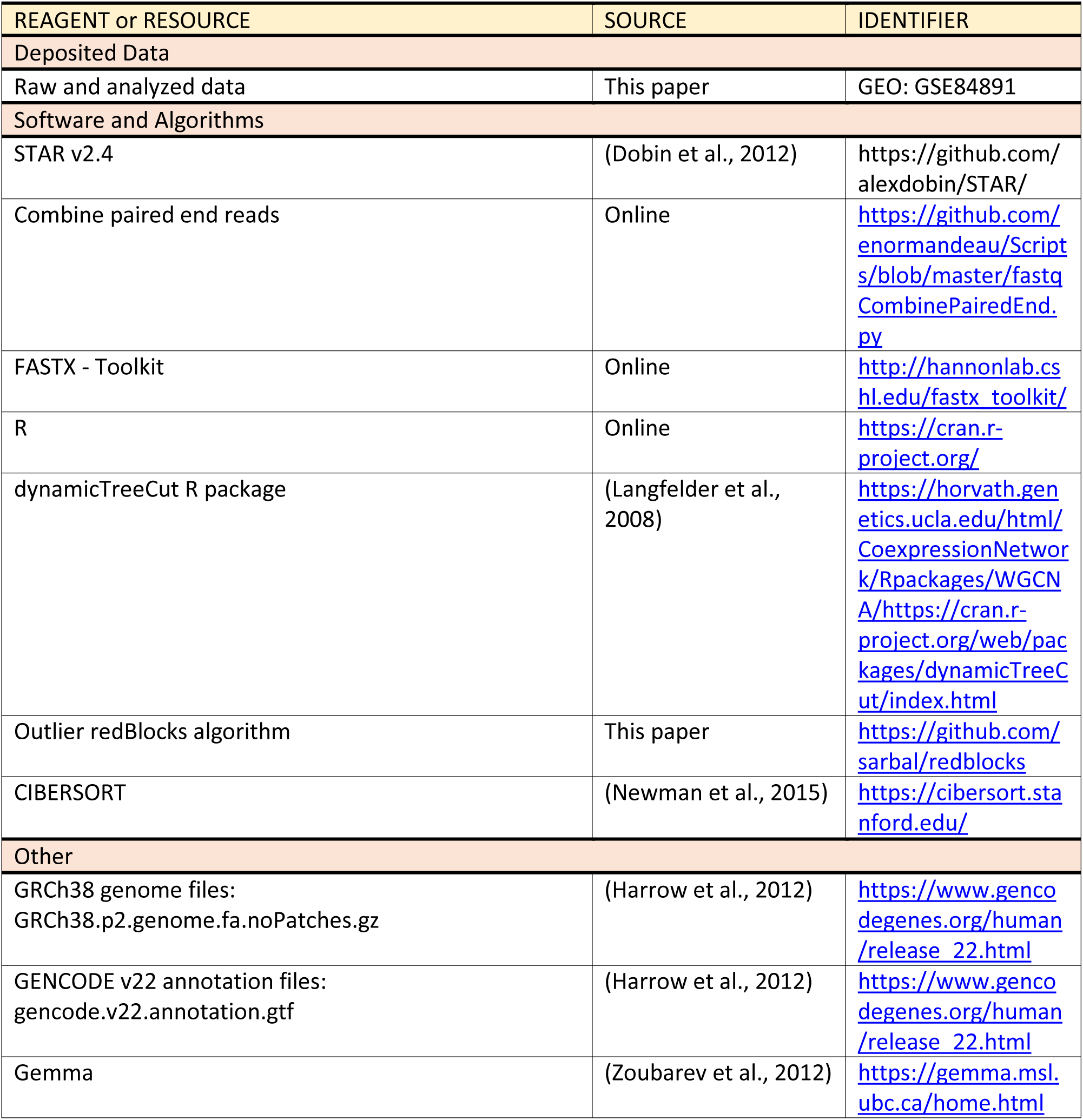

## Supplementary

**Figure S1 Meta-analytic co-expression frequency network generation. Related to Figure 2.**

**Figure S2 ERCC spike-ins QC. Related to Figure 3.**

**Figure S3 Pathway recurrence and GO group size. Related to Figure 4.**

**Figure S4 Robustness assessment of DE threshold. Related to Figure 3.**

**Table S1 Library numbers and adapter sequences used in this study. Related to Figure 3.**

**Table S2 RNA-seq experiments used. Related to Figures 3-6.**

**Table S3 Studies used in the meta-analysis. Related to Figure 6.**

**Table S4 *TAF1* syndrome recurrent genes. Related to Figure 5.**

**Table S5 *TAF1* syndrome enrichment results. Related to Figure 4.**

**Table S6 Huntington’s disease recurrent genes. Related to Figure 6.**

**Table S7 Parkinson’s disease recurrent genes. Related to Figure 6.**

**Table S8 Parkinson’s disease enrichment results. Related to Figure 6.**

**Table S9 Schizophrenia recurrent genes. Related to Figure 6.**

**Table S10 Schizophrenia enrichment results. Related to Figure 6.**

