## Supplemental File for "Not by systems alone: replicability assessment of disease expression signals"

### Supplementary Figures

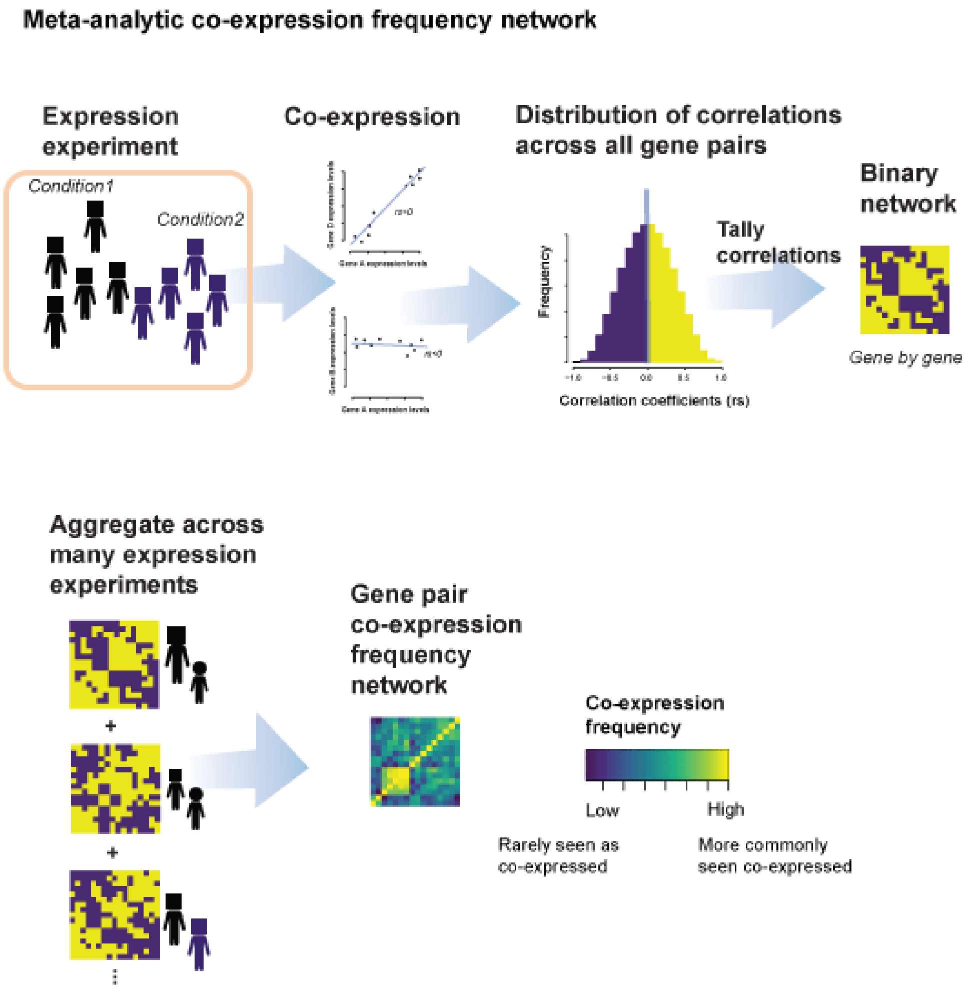

**Figure S1 Meta-analytic co-expression frequency network generation. Related to Figure 2.**
Taking each co-expression distribution, we threshold so that all Spearman's correlation coefficient $r_s>0$ are given a value of 1 (light yellow shading), and all others 0 (purple shading). Repeating for all human RNA-seq experiments, we aggregate these individual networks into an occurrence network, such that for each gene pair we have the frequency these genes share any change in expression across samples. Those closer to 0 are seen less often as co-expressed, while those closer to 1 are commonly seen as co-expressed in other data. These measures are used to define a joint signal i.e., co-functionality/co-regulation of genes.

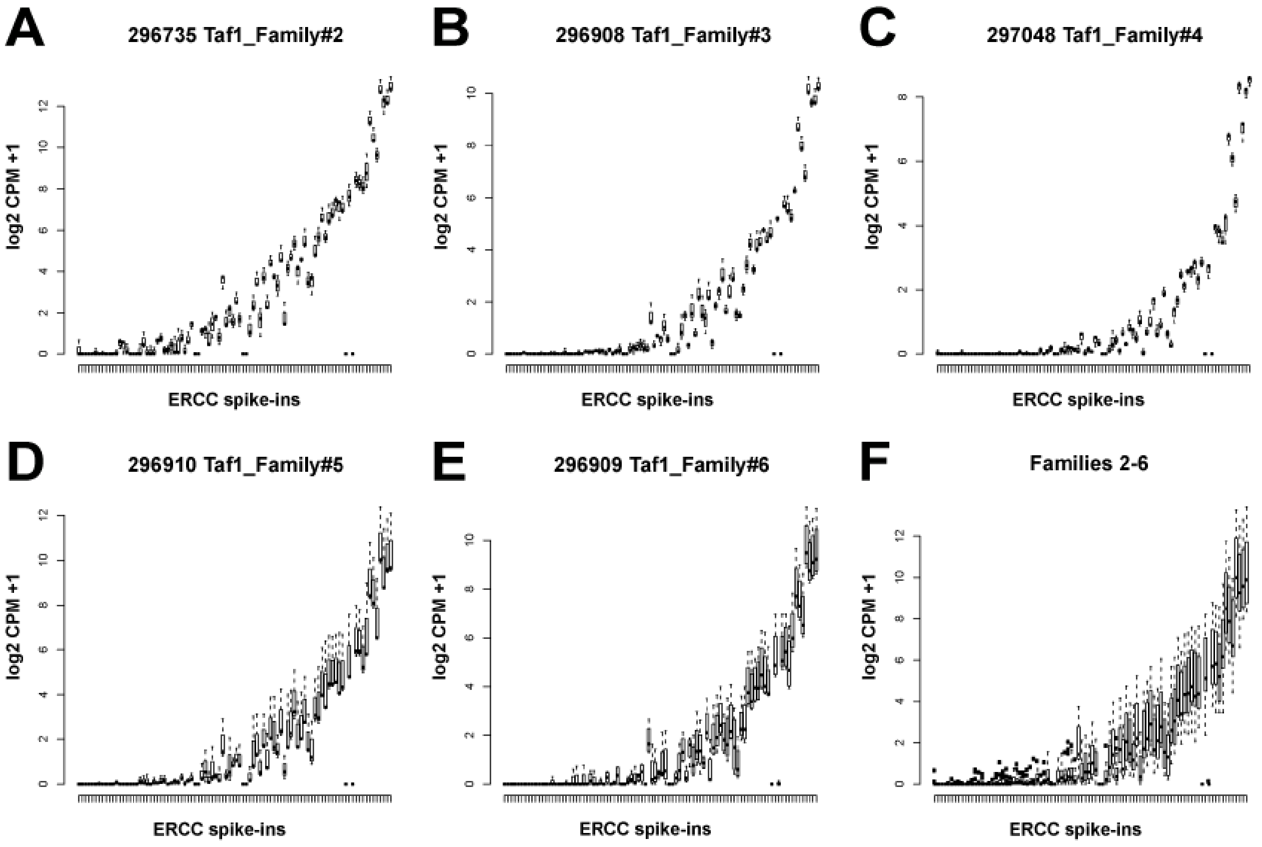

**Figure S2 ERCC spike-ins QC. Related to Figure 3.**
For each sample in each family (batch), we calculate and plot the log2 CPM+1 value of the 92 spike-ins used. The final plot shows the distributions across all families. As Family 1 was sequenced prior, it was not part of this analysis.

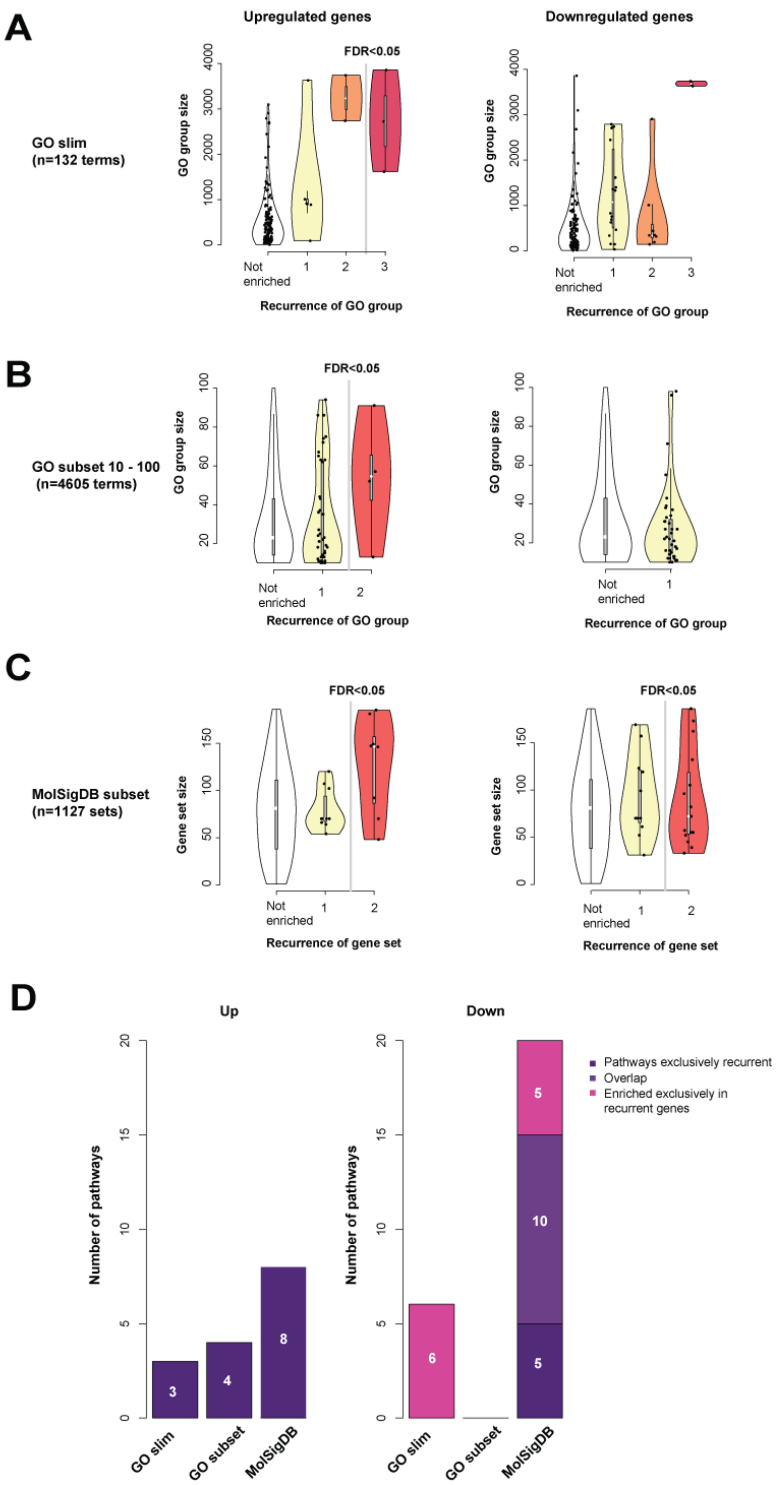

**Figure S3 Pathway recurrence and GO group size. Related to Figure 4.**
(A) Enrichment of GO slim groups compared to recurrence for up- and downregulated genes. (B) Same but for GO terms of gene set size between 10 and 100. (C) MolSigDB enrichment results for HALLMARK, KEGG, BIOCARTA and REACTOME gene sets. (D) Recurrent pathways and their overlap with pathways enriched using only the subset of recurrent genes.

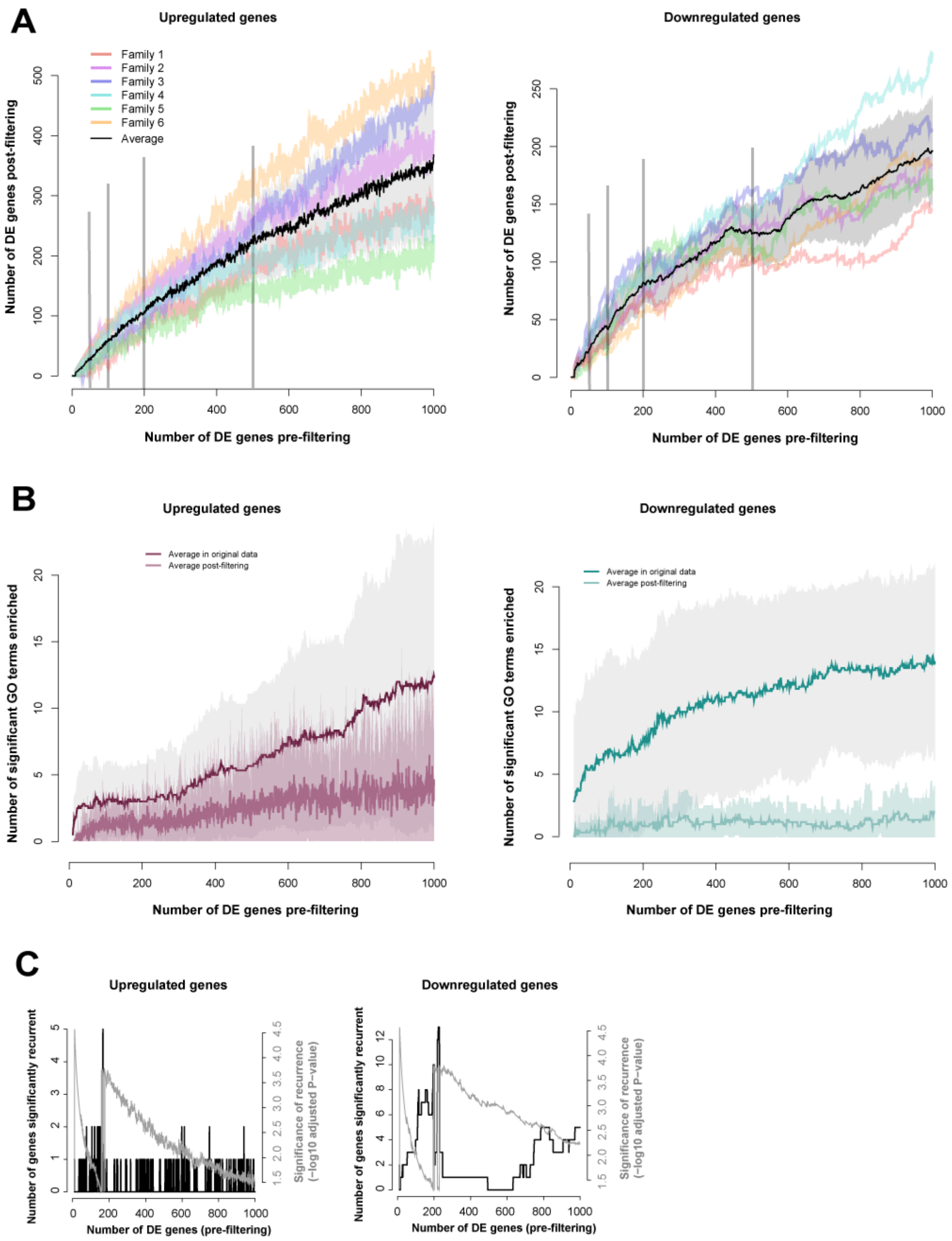

**Figure S4 Robustness assessment of DE threshold. Related to Figure 3.**

(A) Robustness analyses showing the number of genes pre and post co-expression analysis for each family (colored lines) and averaged (black line, SD grey shadow). As we increased the number of genes considered for DE, the number of genes we filter also increases, averaging to approximately two-thirds removed. (B) If we look at the enrichment of these DE gene sets (pre-filtering darker line +/-SD shadow), we see that filtering off the modules removes all but a few significant terms (lighter line, +/-SD shadow). (C) For both increased and decreased expression, the number of significantly recurrent genes post-filtering are few.

### Supplementary Tables

**Table S1 Library numbers and adapter sequences used in this study. Related to Figure 3.**

| | Library | Index | Sequence | SID | Member |
|---|---|---|---|---|---|
| **Family 2** | Taf1_11 | AR002 | CGATGT | 296732 | Proband |
| *Family 6A* | Taf1_12 | AR007 | CAGATC | 296733 | Mother |
| | Taf1_13 | AR019 | GTGAAA | 296734 | Father |
| **Family 3** | Taf1_14 | AR005 | ACAGTG | 296898 | Proband |
| *Family 2A* | Taf1_15 | AR006 | GCCAAT | 296899 | Father |
| | Taf1_16 | AR015 | ATGTCA | 296900 | Mother |
| **Family 4** | Taf1_24 | AR008 | ACTTGA | 297045 | Proband |
| *Family 4A* | Taf1_25 | AR011 | GGCTAC | 297046 | Mother |
| | Taf1_26 | AR022 | CGTACG | 297047 | Father |
| **Family 5** | Taf1_21 | AR001 | ATCACG | 296905 | Proband |
| *Family 5A* | Taf1_22 | AR010 | TAGCTT | 296906 | Father |
| | Taf1_23 | AR020 | GTGGCC | 296907 | Mother* |
| **Family 6** | Taf1_17 | AR003 | TTAGGC | 296901 | Proband |
| *Family 10A* | Taf1_18 | AR009 | GATCAG | 296902 | Father |
| | Taf1_19 | AR022 | CGTACG | 296903 | Mother* |
| | Taf1_20 | AR027 | ATTCCT | 296904 | Sibling |

* Carrier of same mutation. We have relabeled the family IDs from the O'Rawe study (in italics below). Family 1 is the same between studies.

**Table S2** **RNA-seq experiments used. Related to Figures 3 -6.**

| GSE ID | Number of samples | Genes detected | PMID | GSE ID | Number of samples | Genes detected | PMID |
|---|---|---|---|---|---|---|---|
| **GSE12946** | 16 | 52025 | 18978772 | **GSE53239** | 22 | 47936 | 24393808 |
| **GSE22260** | 30 | 45951 | 21571633 | **GSE53450** | 10 | 41211 | NA |
| **GSE25599** | 20 | 55786 | NA | **GSE53635** | 15 | 42921 | NA |
| **GSE30198** | 11 | 33391 | 22454233 | **GSE53949** | 10 | 40902 | 24379348 |
| **GSE32874** | 20 | 40117 | 22424236 | **GSE54456** | 174 | 56284 | 24441097 |
| **GSE33587** | 13 | 11784 | 22922032 | **GSE55504** | 28 | 42767 | NA |
| **GSE33601** | 10 | 33866 | 23029578 | **GSE55758** | 16 | 42014 | NA |
| **GSE34736** | 40 | 35824 | 25358733 | **GSE56087** | 18 | 39702 | 25239642 |
| **GSE35296** | 10 | 41170 | 22412385 | **GSE56267** | 13 | 50129 | NA |
| **GSE36761** | 29 | 50410 | 24459294 | **GSE56787** | 38 | 51441 | 24861551 |
| **GSE39170** | 15 | 56647 | 22931923 | **GSE57148** | 189 | 44547 | NA |
| **GSE40419** | 164 | 50922 | 22975805 | **GSE57253** | 26 | 48780 | NA |
| **GSE43520** | 12 | 50530 | NA | **GSE57369** | 14 | 40925 | NA |
| **GSE43834** | 11 | 38537 | NA | **GSE57945** | 359 | 60607 | 25003194 |
| **GSE45419** | 32 | 45626 | 24223926 | **GSE57982** | 31 | 53742 | 25083870 |
| **GSE45669** | 16 | 42191 | 23861770 | **GSE58135** | 168 | 57446 | NA |
| **GSE46224** | 40 | 39330 | 24429688 | **GSE58150** | 16 | 30360 | NA |
| **GSE46513** | 15 | 48122 | NA | **GSE58387** | 23 | 31790 | 25016029 |
| **GSE46622** | 12 | 51114 | 23874421 | **GSE58434** | 53 | 41601 | NA |
| **GSE46665** | 25 | 36014 | NA | **GSE58441** | 12 | 34162 | 25358733 |
| **GSE47462** | 72 | 55264 | NA | **GSE58608** | 24 | 34501 | 25016029 |
| **GSE47944** | 84 | 47056 | 24909886 | **GSE58640** | 16 | 30641 | 24920534 |
| **GSE48166** | 30 | 40514 | NA | **GSE60216** | 54 | 52798 | NA |
| **GSE48850** | 11 | 41736 | NA | **GSE60590** | 33 | 35560 | 25484259 |
| **GSE48865** | 274 | 51333 | NA | **GSE60591** | 24 | 34501 | NA |
| **GSE49379** | 30 | 48967 | 24866127 | **GSE61141** | 24 | 28953 | NA |
| **GSE50244** | 89 | 43238 | 25015099 | **GSE61162** | 14 | 42719 | 25822018 |
| **GSE50599** | 10 | 33146 | NA | **GSE62098** | 12 | 47513 | NA |
| **GSE50760** | 54 | 43045 | 25049118 | **GSE63042** | 129 | 50635 | NA |
| **GSE51005** | 12 | 36137 | NA | **GSE63311** | 83 | 47747 | NA |
| **GSE51518** | 20 | 47202 | NA | **GSE63420** | 14 | 45978 | NA |
| **GSE52194** | 20 | 50514 | NA | **GSE63646** | 71 | 55235 | NA |
| **GSE52248** | 18 | 41455 | NA | **GSE63979** | 42 | 52744 | NA |
| **GSE52463** | 15 | 50988 | 24647608 | **GSE64813** | 188 | 45851 | NA |
| **GSE52778** | 16 | 36650 | NA | **GSE65000** | 10 | 58274 | 25721503 |
| **GSE52946** | 20 | 48802 | 25381879 | **GSE65446** | 10 | 36108 | NA |
| **GSE53080** | 185 | 27873 | NA | **GSE66622** | 80 | 47131 | NA |
| **GSE67427** | 89 | 49622 | NA | | | | |

Table S3 Studies used in the meta-analysis. **Related to Figure 6**.

| Disease | Dataset label | Samples | Tissue | Method | GemmaID | GSEID | SRAID/ERAID | PMID |
|---|---|---|---|---|---|---|---|---|
| **HD** | 365_GSE3790 | 404 | Brain | Microarray | 365 | GSE3790 | | 16467349 (Hodges et al., 2006) |
| **HD** | 2684_GSE26927 | 118 | Other | Microarray | 2684 | GSE26927 | | 22864814 (Durrenberger et al., 2012) |
| **HD** | 2771_GSE1751 | 31 | Blood | Microarray | 2771 | GSE1751 | | 16043692 (Borovecki et al., 2005) |
| **HD** | 6533_GSE45516 | 9 | Other | Microarray | 6533 | GSE45516 | | 24296361 (Marchina et al., 2014) |
| **HD** | labadorf | 69 | Brain | RNA-seq | | GSE64810 | | 26636579 (Labadorf et al., 2015) |
| **PD** | pdnc | 59 | Blood | Exon array | | | | 21943286 (Kedmi et al., 2011) |
| **PD** | pdconpdd.pdd_vs_con | 29 | Brain | Microarray | | | | 18649390 (Stamper et al., 2008) |
| **PD** | pdcon_wps | 19 | Brain | Microarray | | | | 19052140 (Simunovic et al., 2009) |
| **PD** | pdcon_sn | 8 | Brain | Microarray | | | | 18462474 (Bossers et al., 2009) |
| **PD** | pd_oc_yc.oc_vs_pd | 22 | Brain | Microarray | | | | 21541762 (Elstner et al., 2011) |
| **PD** | pd_ms | 6 | Brain | Microarray | | | | 16626704 (Vogt et al., 2006) |
| **PD** | pd_exprs_g2019s | 40 | Blood | RNA-seq | | | | 25475535 (Infante et al., 2015) |
| **PD** | pd_stria.m | 37 | Brain | Microarray | | | | 25170892 (Riley et al., 2014) |
| **PD** | pdedger | 24 | Skin | RNA-seq | | GSE54285 | | 28068941 (Planken et al., 2017) |
| **PD** | GSE8397 | 94 | Brain | Microarray | 873 | GSE8397 | | 16344956 (Moran et al., 2006) |
| **PD** | GSE7621 | 25 | Brain | Microarray | 549 | GSE7621 | | 17571925 (Lesnick et al., 2007) |
| **PD** | GSE43490 | 41 | Brain | Microarray | 11614 | GSE43490 | | 25525598 (Corradini et al., 2014) |
| **PD** | GSE28894 | 114 | Brain | Microarray | 3349 | GSE28894 | | |
| **PD** | GSE22491 | 18 | Blood | Microarray | 5670 | GSE22491 | | 20096956 (Mutez et al., 2011) |
| **PD** | GSE20141 | 18 | Brain | Microarray | 1885 | GSE20141 | | 20926834 (Zheng et al., 2010) |
| **SCZ** | 2684_GSE26927 | 118 | Brain | Microarray | 2684 | GSE26927 | | 22864814 (Durrenberger et al., 2012) |
| **SCZ** | 2715_GSE25673 | 24 | IPSC | Microarray | 2715 | GSE25673 | | 21490598 (Brennand et al., 2011) |
| **SCZ** | 8361_GSE53987 | 205 | Brain | Microarray | 8361 | GSE53987 | | |
| **SCZ** | 8487_GSE40102 | 24 | IPSC | Microarray | 8487 | GSE40102 | | 24686136 (Brennand et al., 2015) |
| **SCZ** | wu | 18 | Brain | RNA-seq | | | ERP001304 | 22558445 (Wu et al., 2012) |
| **SCZ** | hwang | 29 | Brain | RNA-seq | | | | 24169640 (Hwang et al., 2013) |
| **SCZ** | fillman | 40 | Brain | RNA-seq | | | | 22869038 (Fillman et al., 2012) |
| **SCZ** | xu | 6 | Blood | RNA-seq | | | | 23282246 (Xu et al., 2012) |
| **SCZ** | sainz | 76 | Blood | RNA-seq | | | | 23164819 (Sainz et al., 2012) |
| **SCZ** | chang | 46 | Brain | RNA-seq | | | SRP102186 | 28809853 (Chang et al., 2017) |

**Table S4** ***TAF1*** **syndrome recurrent genes. Related to Figure 5.**

| ***Upregulated*** | | | |
|---|---|---|---|
| **Gene** | **Recurrence** | **Disjoint recurrence** | **Description** |
| **IGLV1-44* | 3 | 3 | Immunoglobulin Lambda Variable 1-44 |
| *FFAR3* | 3 | 2 | Free Fatty Acid Receptor 3 (GPCR) |
| *RN7SK* | 3 | 1 | 7SK Small Nuclear RNA |
| *ISG15* | 3 | 1 | ISG15 Ubiquitin-Like Modifier (interferon related) |
| ***Downregulated*** | | | |
| **Gene** | **Recurrence** | **Disjoint recurrence** | **Description** |
| **IGFBP3* | 4 | 4 | Insulin Like Growth Factor Binding Protein 3 |
| **CACNA1I* | 4 | 4 | calcium voltage-gated channel subunit alpha1 I |
| **OLFM4* | 3 | 3 | olfactomedin 4 |
| **S100B* | 3 | 3 | S100 calcium binding protein B |
| *CFH* | 3 | 2 | Age-Related Maculopathy Susceptibility 1 , Complement Factor H |
| *RPS7* | 4 | 2 | Ribosomal Protein S7 |
| *PRSS30P* | 3 | 2 | serine protease 30 |
| *C1QA* | 3 | 1 | Complement C1q A Chain |
| *SNRPG* | 3 | 1 | Small Nuclear Ribonucleoprotein Polypeptide G |
| *LSM3* | 3 | 1 | LSM3 Homolog, U6 Small Nuclear RNA And MRNA Degradation Associated |
| *RPS3A* | 3 | 1 | Small Ribosomal Subunit Protein ES1 |
| *RPL7* | 3 | 1 | Large Ribosomal Subunit Protein UL30 |
| *KLRB1* | 3 | 1 | Killer Cell Lectin Like Receptor B1 |
| *KIR2DS4* | 3 | 1 | killer cell immunoglobulin like receptor, two Ig domains and short cytoplasmic tail 4 |

* significantly recurrent and disjoint

**Table S5** ***TAF1*** **syndrome enrichment results. Related to Figure 4.**

| *Upregulated* | | | | |
|---|---|---|---|---|
| **GO ID** | **Recurrence** | **Disjoint recurrence** | **Recurrent gene enrichment (p-adjusted)** | **Description** |
| **GO:0002376** | 3 | 3 | 0.986 | immune system process |
| **GO:0006950** | 3 | 3 | 1.000 | response to stress |
| **GO:0007165** | 3 | 3 | 1.000 | signal transduction |
| ***Downregulated*** | | | 0.986 | |
| **GO ID** | **Recurrence** | **Disjoint recurrence** | **Recurrent gene enrichment (p-adjusted)** | **Description** |
| **GO:0003723** | 1 | 1 | 0.035 | RNA binding |
| **GO:0005576** | 3 | 3 | 0.035 | extracellular region |
| **GO:0005840** | 2 | 2 | 0.018 | ribosome |
| **GO:0006412** | 2 | 2 | 0.035 | translation |
| **GO:0006605** | 2 | 1 | 0.035 | protein targeting |
| **GO:0034655** | 2 | 1 | 0.007 | nucleobase-containing compound catabolic process |

**Table S6 Huntington's disease recurrent genes. Related to Figure 6.**

| ***Upregulated*** | | | |
|---|---|---|---|
| **Gene** | **Recurrence** | **Disjoint recurrence** | **Description** |
| **TXNIP* | 3 | 3 | Thioredoxin Interacting Protein |
| **PRKX* | 3 | 3 | Protein Kinase, X-Linked |
| **ANG* | 3 | 3 | Angiogenin |
| **GMPR* | 3 | 3 | Guanosine Monophosphate Reductase |
| **PLA1A* | 3 | 3 | Phospholipase A1 Member A |
| *DDIT4* | 3 | 2 | DNA Damage Inducible Transcript 4 |
| *KCNE4* | 3 | 2 | Potassium Voltage-Gated Channel Subfamily E Regulatory Subunit 4 |
| *MT1F* | 3 | 2 | Metallothionein 1F |
| *SLC14A1* | 3 | 2 | Solute Carrier Family 14 Member 1 (Kidd Blood Group) |
| *SLC16A9* | 3 | 2 | Solute Carrier Family 16 Member 9 |
| *ANGPT1* | 4 | 1 | Angiopoietin 1 |
| *EMP1* | 3 | 1 | Epithelial Membrane Protein 1 |
| *EMP3* | 3 | 1 | Epithelial Membrane Protein 3 |
| *ID3* | 3 | 1 | Inhibitor Of DNA Binding 3, HLH Protein |
| *MTHFD2* | 3 | 1 | Methylenetetrahydrofolate Dehydrogenase (NADP+ Dependent) 2, Methenyltetrahydrofolate Cyclohydrolase |
| *CLEC2B* | 3 | 0 | C-Type (Calcium Dependent, Carbohydrate-Recognition Domain) Lectin |
| *SCGB1D2* | 3 | 0 | Secretoglobin Family 1D Member 2 |
| *UHRF1* | 3 | 0 | Ubiquitin Like With PHD And Ring Finger Domains 1 |
| ***Downregulated*** | | | |
| **Gene** | **Recurrence** | **Disjoint recurrence** | **Description** |
| *CACNA1E* | 3 | 2 | another calcium channel |
| *SV2C* | 3 | 1 | synaptic vesicle glycoprotein |
| *IMPG1* | 3 | 2 | retinal interphotoreceptor matrix glycoprotein |
| *NRGN* | 3 | 1 | Neurogranin (postsynaptic protein kinase substrate) |
| *YPEL1* | 3 | 0 | Yippee Like 1 |
| *HTR2C* | 3 | 2 | 5-Hydroxytryptamine Receptor 2C (responds to signaling through the neurotransmitter serotonin, depression candidate gene) |

* significantly recurrent and disjoint

**Table S7 Parkinson's disease recurrent genes. Related to Figure 6.**

| ***Upregulated*** | | | |
|---|---|---|---|
| **Gene** | **Recurrence** | **Disjoint recurrence** | **Description** |
| *GPR88* | 3 | 3 | G Protein-Coupled Receptor 88 ( protein-coupled receptor found almost exclusively in the striatum; defects associated with learning difficulties and neuropsychiatric disorders) |
| *NDST3* | 3 | 3 | N-Deacetylase And N-Sulfotransferase 3 |
| *ANKRD37* | 3 | 3 | Ankyrin Repeat Domain 37 |
| *HSPA1A* | 3 | 3 | Heat Shock Protein Family A (Hsp70) Member 1A |
| *CA2* | 3 | 3 | Carbonic Anhydrase 2 |
| *THY1* | 3 | 3 | Thy-1 Cell Surface Antigen (May play a role in cell-cell or cell-ligand interactions during synaptogenesis and other events in the brain.) |
| *CIT* | 4 | 3 | Citron Rho-Interacting Serine/Threonine Kinase (central nervous system development) |
| *NUPR1* | 3 | 3 | Nuclear Protein 1, Transcriptional Regulator |
| *CBR1* | 3 | 3 | Carbonyl Reductase 1 |
| *MAFF* | 3 | 3 | MAF BZIP Transcription Factor F |
| ***Downregulated*** | | | |
| **Gene** | **Recurrence** | **Disjoint recurrence** | **Description** |
| **SNCA* | 6 | 6 | Synuclein Alpha (known parkinsons disease) |
| **UCHL1* | 5 | 5 | Ubiquitin C-Terminal Hydrolase L1 (associated with parkinsons) |
| **SLC10A4* | 4 | 4 | Solute Carrier Family 10 Member 4 |
| **DLK1* | 4 | 4 | Delta Like Non-Canonical Notch Ligand 1 |
| *MLLT11* | 4 | 3 | MLLT11, Transcription Factor 7 Cofactor |
| *GAP43* | 4 | 3 | Growth Associated Protein 43 (axonal regeneration) |
| *HBB* | 4 | 3 | Hemoglobin Subunit Beta |
| *MOAP1* | 4 | 3 | Modulator Of Apoptosis 1 |
| *NMNAT2* | 4 | 2 | Nicotinamide Nucleotide Adenylyltransferase 2 |
| *NR4A2* | 4 | 2 | Nuclear Receptor Subfamily 4 Group A Member 2 (associated with parkinsons) |
| *SCN3A* | 4 | 2 | Sodium Voltage-Gated Channel Alpha Subunit 3 |
| *OSBPL10* | 4 | 2 | Oxysterol Binding Protein Like 10 |
| *SNX10* | 4 | 2 | Sorting Nexin 10 |
| *SLC18A2* | 5 | 2 | Solute Carrier Family 18 Member A2 (synaptic vesicle function) |
| *CMAS* | 4 | 2 | Cytidine Monophosphate N-Acetylneuraminic Acid Synthetase |
| *SYT1* | 5 | 2 | Synaptotagmin 1 |
| *CIRBP* | 4 | 2 | Cold Inducible RNA Binding Protein |
| *AKAP12* | 4 | 1 | A-Kinase Anchoring Protein 12 |
| *AMPH* | 4 | 1 | Amphiphysin ( cytoplasmic surface of synaptic vesicles) |
| *HPRT1* | 5 | 1 | Hypoxanthine Phosphoribosyltransferase 1 |
| *DMXL2* | 4 | 0 | Dmx Like 2 |

* significantly recurrent and disjoint

**Table S8 Parkinson's disease enrichment results. Related to Figure 6.**

| *Upregulated* | | | | |
|---|---|---|---|---|
| **GO ID** | **Recurrence** | **Disjoint recurrence** | **Recurrent gene enrichment (p-adjusted)** | **Description** |
| **GO:0007267** | 5 | 5 | 1.000 | cell-cell signaling |
| **GO:0030154** | 5 | 5 | 0.515 | cell differentiation |
| **GO:0040011** | 5 | 5 | 1.000 | locomotion |
| **GO:0048856** | 5 | 5 | 0.515 | anatomical structure development |
| **Downregulated** | | | | |
| **GO IDGene** | **Recurrence** | **Disjoint recurrence** | **Recurrent gene enrichment (p-adjusted)** | **Description** |
| **GO:0005886** | 6 | 5 | 0.169 | plasma membrane |
| **GO:0007267** | 6 | 6 | 0.169 | cell-cell signaling |
| **GO:0008289** | 2 | 1 | 0.006 | lipid binding |
| **GO:0016023** | 6 | 3 | 0.006 | cytoplasmic membrane-bounded vesicle |
| **GO:0016192** | 6 | 5 | 0.074 | vesicle-mediated transport |

**Table S9 Schizophrenia recurrent genes. Related to Figure 6.**

| *Upregulated* | | | |
|---|---|---|---|
| **Gene** | **Recurrence** | **Disjoint recurrence** | **Description** |
| **FCN3* | 4 | 3 | Ficolin 3 (lectin pathways, SLE, immunity related) |
| *CHI3L2* | 3 | 0 | Chitinase 3 Like 2 |
| *ADAMTS9* | 3 | 2 | ADAM Metallopeptidase With Thrombospondin Type 1 Motif 9 |
| *IFITM2* | 3 | 1 | Interferon Induced Transmembrane Protein 2 |
| *IFITM1* | 3 | 0 | Interferon Induced Transmembrane Protein 1 |
| *TNFSF14* | 3 | 1 | TNF Superfamily Member 14 |
| *KLK3* | 3 | 2 | Kallikrein Related Peptidase 3 |
| *SLCO4A1* | 3 | 1 | Solute Carrier Organic Anion Transporter Family Member 4A1 |
| ***Downregulated*** | | | |
| **Gene** | **Recurrence** | **Disjoint recurrence** | **Description** |
| **PPP1R17* | 3 | 3 | Protein Phosphatase 1 Regulatory Subunit 17 (hypercholesterolemia) |
| *SPP1* | 3 | 2 | Secreted Phosphoprotein 1 (cytokine that upregulates expression of interferon-gamma and interleukin-12) |
| *OLR1* | 3 | 2 | Oxidized Low Density Lipoprotein Receptor 1( may modify the risk of Alzheimer's disease) |
| *SPARCL1* | 3 | 1 | SPARC Like 1 (calcium ion binding) |

* significantly recurrent and disjoint

**Table S10 Schizophrenia enrichment results. Related to Figure 6.**

| *Upregulated* | | | | |
|---|---|---|---|---|
| **GO ID** | **Recurrence** | **Disjoint recurrence** | **Recurrent gene enrichment (p-adjusted)** | **Description** |
| **GO:0002376** | 2 | 1 | 0.031 | immune system process |
| **GO:0005576** | 4 | 4 | 0.964 | extracellular region |
| **GO:0005615** | 4 | 4 | 0.514 | extracellular space |
| ***Downregulated*** | | | | |
| **GO ID** | **Recurrence** | **Disjoint recurrence** | **Recurrent gene enrichment (p-adjusted)** | **Description** |
| **GO:0005615** | 3 | 2 | 1.000 | extracellular space |
| **GO:0007267** | 3 | 3 | 1.000 | cell-cell signaling |
